# Spatial suppression and sensitivity for motion in schizophrenia

**DOI:** 10.1101/799395

**Authors:** Daniel Linares, Silvia Amoretti, Rafael Marin-Campos, André Sousa, Laia Prades, Josep Dalmau, Miquel Bernardo, Albert Compte

## Abstract

Perceptual spatial suppression is a phenomenon in which the perceived strength of a stimulus in some region of the space is reduced when the stimulus is surrounded by other stimuli. For contrast perception, several studies suggest that spatial suppression is reduced in patients with schizophrenia. For motion perception, only one study has been conducted in a cohort of 16 patients, suggesting that spatial suppression is reduced. It is unknown, however, whether this reduction is related to the lower IQ that schizophrenic patients usually show; as there is evidence that spatial suppression for motion increases with IQ in healthy individuals. Here, we sought to determine the spatial suppression for motion in a larger cohort of 33 patients with schizophrenia controlling for IQ. We found a weakened spatial suppression in patients with schizophrenia, consistent with the previous study (g = 0.47, CI = [0.055, 0.88], combining the previous and our study). For comparison, we performed a meta-analysis on spatial suppression for contrast and found a similar effect size. We found that patients had a lower IQ than controls, but this difference did not explain their weaker spatial suppression. Further, we found that spatial suppression of patients, but not controls increased with their IQ and decreased with age in both groups. Finally, as we estimated lapses of attention, we could estimate motion sensitivity and found that it was decreased in patients. We speculate about possible alterations in neurotransmission that might explain the reduced spatial suppression and sensitivity that we found.

## INTRODUCTION

Hallucinations, perceptual experiences that occur without stimuli, are a defining feature of schizophrenia (American Psychiatric Association 2013). In addition to these powerful perceptual anomalies, there is evidence that stimulus perception is also affected (Butler, Silverstein, and Dakin 2008; Javitt and Sweet 2015; Yoon et al. 2013; Y. Chen 2011). For example, it has been suggested that perceptual spatial suppression is reduced in patients with schizophrenia (Tadin et al. 2006; Dakin, Carlin, and Hemsley 2005). Perceptual spatial suppression is a perceptual phenomenon in which the perceived strength of a stimulus in some region of the space is reduced when the stimulus is surrounded by other stimuli. The phenomenon is linked to gain control adjustments (Carandini and Heeger 2011; Butler, Silverstein, and Dakin 2008) and the segmentation of objects from their background (Allman, Miezin, and McGuinness 1985; Tadin et al. 2019). A reduced spatial suppression in schizophrenia, thus, should be associated with an impairment on those fundamental visual functions.

Two pieces of evidence suggest that the reduction of spatial suppression in schizophrenia is likely a genuine perceptual alteration rather than a generalized behavioural deficit, such as a lack of attention or motivation (Skottun and Skoyles 2007; Yoon et al. 2013). First, a reduced spatial suppression can result in patients performing the perceptual task more accurately than healthy individuals (Tadin et al. 2006; Dakin, Carlin, and Hemsley 2005). Second, spatial suppression is often characterized as the difference in perceptual sensitivity with and without a surround stimulus, and thus is not affected by a global change in sensitivity (Tadin et al. 2006; Yoon et al. 2013).

Spatial suppression has been assessed for different perceptual attributes (Yang et al. 2013; Tibber et al. 2013). For contrast, Dakin and colleagues (Dakin, Carlin, and Hemsley 2005), reported that patients with schizophrenia showed a strongly reduced perceptual spatial suppression. The spatial suppression for contrast, often called the contrast-contrast effect, describes the reduction in apparent contrast of a central stimulus that occurs when it is surrounded by a high contrast stimulus (Chubb, Sperling, and Solomon 1989). Later studies have found evidence consistent with this alteration, although of weaker magnitude (Barch et al. 2012; Yang et al. 2013; Tibber et al. 2013; Serrano-Pedraza et al. 2014; Yoon et al. 2009; M-P Schallmo, Sponheim, and Olman 2015; see also Mannion, Donkin, and Whitford 2017).

For the perception of motion, Tadin and colleagues (Tadin et al. 2006) reported that patients with schizophrenia show a reduced perceptual spatial suppression. Spatial suppression for motion describes the reduction in apparent motion strength of a high contrast central stimulus when it is surrounded by a high contrast stimulus moving in the same direction (Tadin, Lappin, and Blake 2006; Neri and Levi 2009) or the related phenomenon by which the sensitivity to discriminate the motion direction of a high contrast stimulus decreases as its size increases (Tadin et al. 2003). Recent evidence from studies in healthy individuals has suggested that spatial suppression for motion increases with intelligence quotient (IQ) (Arranz-Paraíso and Serrano-Pedraza 2018; Melnick et al. 2013; but see Troche et al. 2018). Given that patients with schizophrenia usually have lower IQ than healthy participants (Van Haren et al. 2019), the reported weaker spatial suppression in patients with schizophrenia (Tadin et al. 2006) could be a result of the patients’ lower IQ. Here, we aimed at replicating spatial suppression for motion in a larger cohort than the previous study (Tadin et al. 2006), while controlling for IQ, and compare the magnitude of the effect with that of spatial suppression for contrast.

## METHODS

### Participants

The study was approved by the ethical committee of the Hospital Clinic of Barcelona and followed the requirements of the Helsinki convention. All participants reported normal or corrected-to-normal visual acuity, did not know the hypothesis of the experiment and provided informed consent.

We recruited 37 outpatients diagnosed with schizophrenia according to DSM-V (American Psychiatric Association 2013) and 33 healthy controls. Patients were between 18 and 65 years old at the time of the first evaluation. Exclusion criteria for both groups were: intellectual disability according to DSM-V criteria, a history of head trauma with loss of consciousness or an organic disease with mental repercussions. For healthy controls, exclusion criteria also included having a first degree relative with a history of psychotic disorder, current or past diagnosis of a psychotic disorder, major depression or other serious psychiatric illnesses such as bipolar disorder.

For 4 patients and 2 controls, motion sensitivity could not be estimated: the accuracy was at or below chance level indicating that the participants responded left for rightward moving stimuli and vice versa. These participants were not included in the analysis.

### Perceptual test

The test was performed in a room with normal fluorescent lighting on a tablet (iPad 2017; 9.7 inches, 2048×1536 pixels, GPU PowerVR GT7600) that we have previously validated for the task used in this study (Linares et al. 2018).

The stimuli (illustrated in Figure 1A) were sinusoidal gratings (0.42 Michelson contrast) of 1 cycle per deg (of visual angle) drifting at 4 deg/s with a Gaussian envelope with a standard deviation of 0.5 deg for the small grating and 2 deg for the large grating. This envelope results in visible stimulus sizes of about 1 and 4 deg. The gratings were displayed in the center of the screen. On each trial, their initial phase was chosen randomly from a range of 5 values (0, 72, 144, 216 and 288°). The background luminance was 32 cd/m^2^.

**Figure 1.**
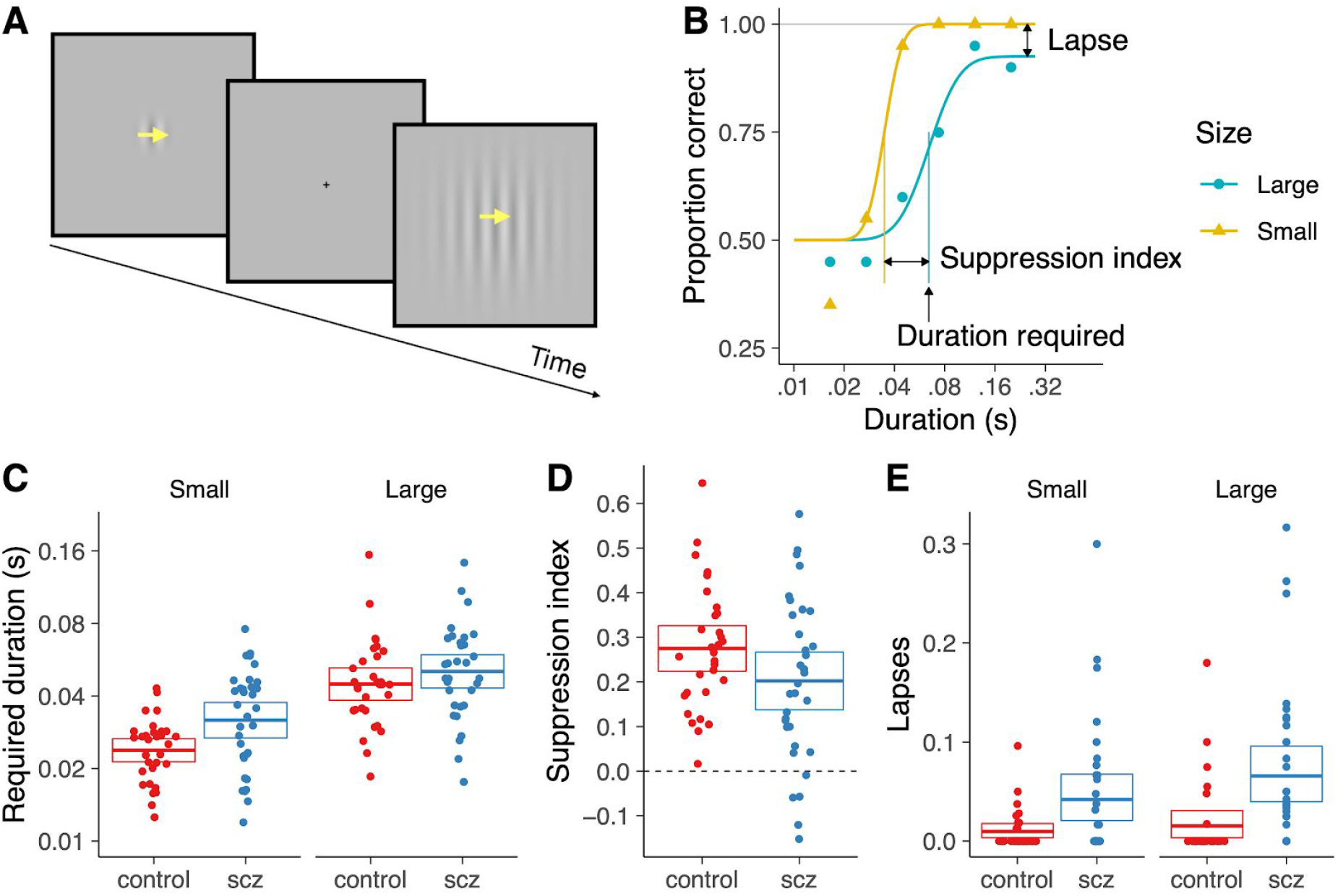
(A) Illustration of the perceptual task. (B) Psychometric function model fitted to one example participant. (C) Motion sensitivity for small and large stimuli for all participants (scz: *patients* with schizophrenia, control: healthy controls). The dots show the results for each participant. The boxes indicate the mean and the 95% confidence intervals. (D). Suppression index for all participants. (E) Lapses for all participants.

To set up the viewing distance, at the beginning of each block the experimenter used a ruler to measure the distance from the eyes to the screen and asked the participant to change position until the distance was about 57 cm. Once the participant told the experimenter that she was in a comfortable position, we asked her to hold that position for the whole block. The experimenter was in the same experimental room controlling that the participant did not change position. Participants were asked to look at the center of the screen during the entire test.

Each trial (Figure 1A) started with the presentation of a cross for 0.3 s. Then, a grating moving to the left or to the right (chosen at random on each trial) was presented. The duration of the grating was controlled using a temporal Gaussian envelope for the contrast of the grating and was chosen at random on each trial from a range of 6 logarithmically spaced durations between 0.02 s and 0.2 s. These durations defined 2 standard deviations of the temporal Gaussian envelope. The peak of the Gaussian envelope occurred 0.3 s after the offset of the cross. For some participants (14 patients and 1 control), we also presented a nominal duration of 0.01 s that we did not consider for the analysis presented here because the duration was not displayed appropriately as described in our previous study (Linares et al. 2018). Nevertheless, when those durations were included the impact on the estimated parameters was negligible.

Participants reported the perceived motion direction by tapping on the left or right part of the screen. We informed participants that they did not need to respond quickly. Feedback was not provided. The next trial started 0.3 s after the response.

The test consisted of 240 trials (6 durations x 2 sizes x 2 directions x 5 initial phases x 2 repetitions). Before the test, participants performed a few training trials with long duration stimuli.

### Clinical, neuropsychological and sociodemographic data

The clinical, neuropsychological and sociodemographic data are presented in Table 1.

**Table 1.** Clinical, neuropsychological and sociodemographic data of the participants. The p-value was calculated using t-test for all comparisons except for the “Number of males” comparison, for which a Chi-squared test was applied.

|  | Controls (n = 31) | Patients (n = 33) | p-value |
| --- | --- | --- | --- |
| Number of males | 21 | 24 | 0.871 |
| Age (years) | 38.6 (SD 13.8) | 38.8 (SD 10.4) | 0.954 |
| IQ | 109 (SD 11) | 102 (SD 12.6) | 0.0128 |
| SES | 3.45 (SD 0.961) | 3.7 (SD 1.21) | 0.371 |
| GAF | 87.5 (SD 6.86) | 63.3 (SD 9.97) | 2.44e-16 |
| PANSS-NEGATIVE | - | 15.3 (SD 5.23) | - |
| PANSS-POSITIVE | - | 9.76 (SD 2.92) | - |
| PANSS-GENERAL | - | 25.7 (SD 6.58) | - |
| PANSS-TOTAL | - | 50.5 (SD 12.1) | - |
| BNSS | - | 15.3 (SD 8.18) | - |
| CPZ EQUIVALENT (mg/day) | - | 382 (SD 282) | - |

The age range was from 19 to 61 years (interquartile range: 13 years) in patients with schizophrenia and from 18 to 67 years (interquartile range: 17 years) in healthy participants.

Psychopathological assessment was carried out with the Spanish validated versions of the Positive and Negative Syndrome Scale (PANNS; Kay, Fiszbein, and Opler 1987; Peralta and Cuesta 1994) and the Brief Negative Symptoms Scale (Mané et al. 2014; BNSS; Kirkpatrick et al. 2011). Higher scores indicate greater severity.

The overall functional outcome was assessed by the Global Assessment of Functioning (GAF; DSM-IV 2010). The GAF is a scale designed to assess the severity of symptoms and the level of functioning. Higher scores correspond to better functioning.

A general IQ composite index was derived from the Vocabulary and Block Design subtest scores of the Wechsler Adult Intelligence Scale for adults (WAIS-III; Wechsler 1997).

The pharmacological treatment was measured by chlorpromazine equivalents (CPZ). Habits of drug abuse were assessed using an adapted version of the European Adaptation of a Multidimensional Assessment Instrument for Drug and Alcohol Dependence scale (Kokkevi and Hartgers 1995).

Education and socioeconomic status (SES) were determined using Hollingshead’s Two-Factor Index of Social Position (Hollingshead and Redlich 1958).

Patients were matched with healthy controls in age, gender, and SES (Table 1).

### Meta-analysis

To find studies that measured contrast spatial suppression in schizophrenia we used Google Scholar. First, we looked up the studies that cited the paper of Dakin and colleagues (Dakin, Carlin, and Hemsley 2005) and identified six studies (Yoon et al. 2009; Barch et al. 2012; Serrano-Pedraza et al. 2014; Yang et al. 2013; Tibber et al. 2013; M-P Schallmo, Sponheim, and Olman 2015). Second, we looked up the references cited in those six studies as well as the studies that cited these six studies and did not identify any additional studies. Third, we searched for the keywords contrast, suppression and schizophrenia. We also did not identify any additional studies.

To find studies that measured motion spatial suppression in schizophrenia we also used Google Scholar. First, we looked up the studies that cited the paper of Tadin and colleagues (Tadin et al. 2006) and did not identify any study. Second, we searched for the keywords motion, suppression and schizophrenia. We also did not retrieve any additional studies.

To perform the meta-analyses, we contacted the authors of the studies, who sent us the anonymized perceptual measures for each participant or the necessary statistics to calculate the effect size and its standard error. As a measure of effect size, we use Hedge’s *g.* For each study, we calculated g and its standard error using the R packages *esc* (Lüdecke 2018) and *meta* (Schwarzer and Others 2007). We used a random-effects model meta-analysis.

### Data analysis

For the perceptual test, we used the R package *quickpsy* (Linares and López i Moliner 2016) to fit the following 3-parameter psychometric function model (Kingdom and Prins 2016):

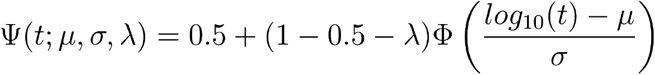

where *t* is the duration of the stimulus, *μ* (motion sensitivity) corresponds to the required duration to respond correctly in about 75% of the trials (the exact proportion is 0.75 – 0.5λ), *σ* corresponds to the slope of the psychometric function, λ is the lapse rate, and Φ is the cumulative normal function.

To assess how motion sensitivity was distorted when lapses were not taken into account, we also fitted the model without including lapse rate—in fact, we included a very small fixed lapse rate (λ = 0.01) to minimize bias in the estimation of motion sensitivity (Prins 2012).

The bootstrap confidence intervals (percentile method) and the permutation tests used 30,000 samples. The performed t-tests were Welch t-tests and the correlations were Pearson correlations.

## RESULTS

### Motion sensitivity and spatial suppression

On each trial (Figure 1A), a briefly-presented grating (small or large) drifted leftwards or rightwards (the direction was chosen at random with equal probability) and the participant reported the perceived direction. Figure 1B shows, for one example participant, the proportion of correct direction discriminations for the two sizes of the stimulus as a function of its duration. For each size, we fitted a psychometric function model to the proportion of correct responses with respect to stimulus duration (see Methods). The model includes a parameter (*μ* of the cumulative normal function) that is related to the duration required to respond correctly in a given proportion of the trials. This is a measure of motion sensitivity: short required durations indicate high sensitivity. The model also includes a lapse parameter (Kingdom and Prins 2016) related to the proportion of incorrect responses for stimuli that should be easy to discriminate as they have been presented for long durations. Incorporating lapses is important because it allows assessing sensitivity independently of the inattentiveness or blinks that might occur in some trials (Barch et al. 2012; Yoon et al. 2013; Dakin, Carlin, and Hemsley 2005; Prins 2013). The required duration and lapse parameters are illustrated in Figure 1B for the model that fits large stimulus trials. For each participant, we also calculated the suppression index (Tadin et al. 2006) as the difference in log units between the required durations for large and small stimuli (Figure 1B).

To assess sensitivity, we used the parameter *μ* instead of the duration threshold, which is another popular measure of performance. The duration threshold is defined as the duration required to respond accurately in a given proportion of trials—75% for example. We used *μ* instead of the duration threshold because the duration threshold depends on lapses and, thus, does not provide a pure measure of motion sensitivity (Prins 2013). If a participant, for example, loses attention in a given proportion of trials, this will increase the duration threshold, but will not affect the sensitivity parameter *μ*. The 75% duration threshold coincides with the value of the sensitivity parameter *μ* in the absence of lapses.

For the required duration to discriminate motion, we performed an ANOVA across participants with size (large or small) as a within-subject factor and group (patient or control) as a between-subjects factor (Figure 1C). Replicating the phenomenon of spatial suppression (Tadin et al. 2003, 2006), motion sensitivity was worse for large stimuli than for small stimuli (F(1,62) = 130, p = 7.6 x 10^-17^; paired g = 1.4, CI = [1.0, 1.8]). In addition, motion sensitivity was worse for patients than controls (F(1,62) = 5.3, p = 0.024; g = 0.43, CI = [0.074, 0.78]), which is consistent with the trend observed by Tadin and colleagues (Tadin et al. 2006).

The previous study (Tadin et al. 2006) reported that compared with healthy participants, the decrease in sensitivity with size (using 3 sizes) was less pronounced in patients with schizophrenia. They reported an interaction of size by group (F(2, 27) = 1.72) with a p=0.19 (Tadin et al. 2006). In our data, the interaction of size by group (F(1, 62) = 3.5) had a p= 0.068. The motion sensitivity of patients was especially impaired for small stimuli (small stimuli: t-test, t(55) = 3.0, p = 0.0038, g = 0.74, CI = [0.22, 1.3]; large stimuli: t-test, t(55) = 1.2, p = 0.24, g = 0.29, CI = [-0.21, 0.79]). To further assess the interaction, we compared the suppression index across groups (Figure 1D). We found that the suppression index was smaller for patients (t-test, t(60) = 1.9; p = 0.066; g = 0.44, CI = [-0.060, 0.94]). Indeed, for 5 patients, we found a negative suppression index, which indicates summation instead of suppression. The reduced suppression in patients is in the direction of the previous study (t-test, t(25) = 1.5; p = 0.14; g = 0.52, CI = [-0.22, 1.3]; statistics calculated using the data shared by Duje Tadin).

For lapses (Figure 1E), as they are proportions we performed permutation tests instead of an ANOVA. Lapses were 3.5 times larger in patients than in controls (the difference in lapse rate was 0.041, this statistic had p = 3.3 x 10^-5^). In addition, lapses for large stimuli were 1.5 times larger than for small stimuli (the difference in lapse rate was 0.015, this statistic had p = 0.0099). The interaction of size by group had a p = 0.12 (the statistic was the difference across groups of participants of the difference in lapses across the two sizes for each participant, which was 0.018).

### Intelligence Quotient (IQ)

In relation to the possible association between spatial suppression and IQ (Arranz-Paraíso and Serrano-Pedraza 2018; Melnick et al. 2013; Troche et al. 2018), we did not find evidence supporting the association in healthy participants, but found evidence for the association in patients (Figure 2A). The interaction of IQ by group (F(1,60) = 4.88) had a p=0.031. The larger spatial suppression concomitant with higher IQ in patients was due to a decrease in motion sensitivity with IQ for large stimuli (Figure 2A).

**Figure 2.**
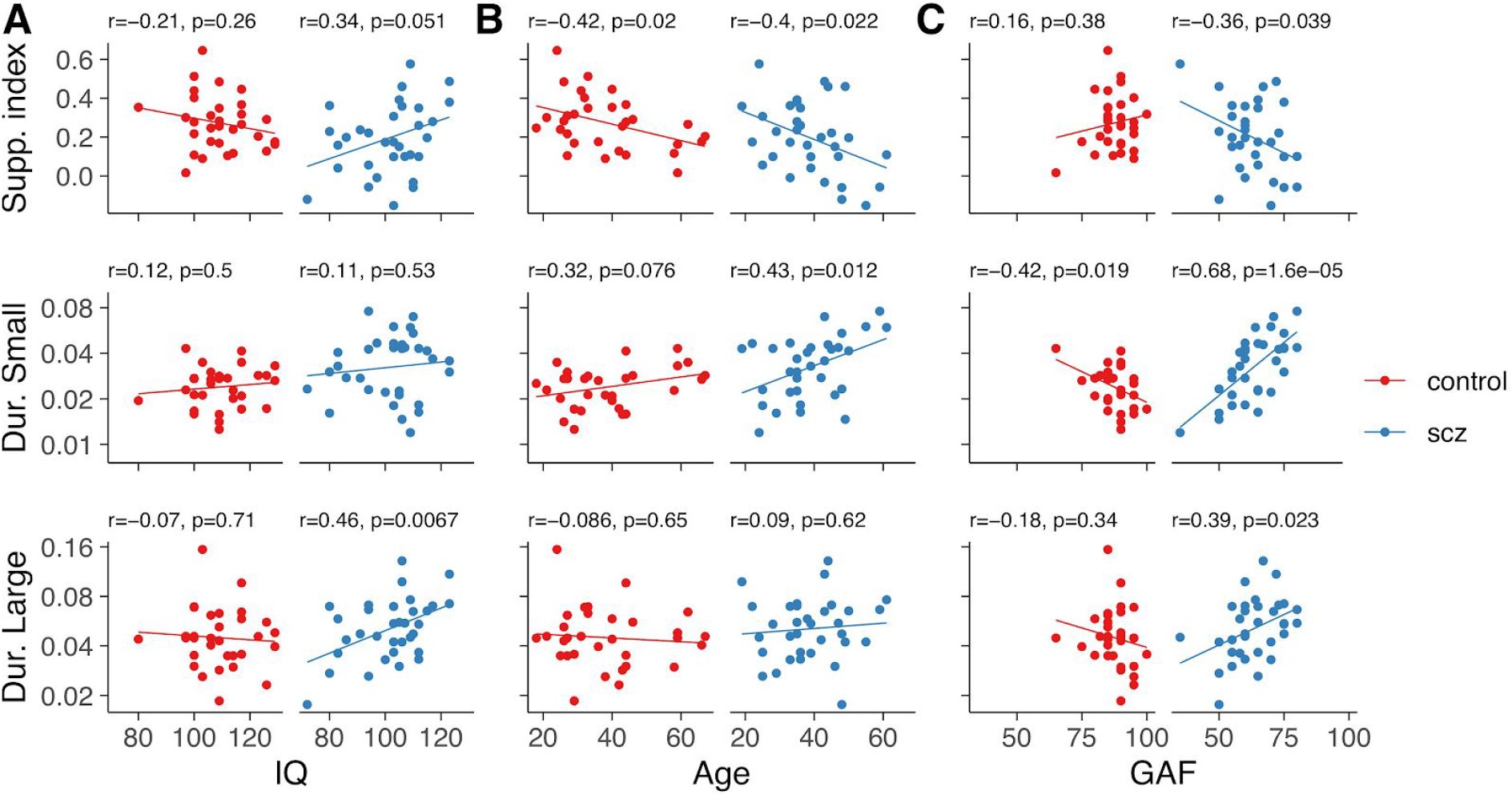
Spatial suppression and motion sensitivity (required durations) in *patients* with schizophrenia (scz) and healthy controls (control) for large and small stimuli against (A) intelligence quotient (IQ), (B) age and (C) the Global Assessment of Functioning.

Patients had lower IQ than controls (Table 1). To assess whether the differences in IQ might explain the differences in spatial suppression, we chose the 27 patients with highest IQ and the 27 controls with lowest IQ. These two groups of participants did not differ in IQ (mean IQ patients: 106, mean IQ controls: 106), age (mean age patients: 39 years, mean age controls: 38 years) or education/socioeconomic status (mean SES patients: 3.5, mean SES controls: 3.5) and we found that the difference in the suppression index across groups was maintained at the same level (t-test, t(49) = 1.7; p = 0.097; g = 0.45, CI = [-0.10, 1.0]). Furthermore, for these two subgroups, as for the whole sample, motion sensitivity was worse for patients than controls (F(1, 52) = 6.7, p = 0.012; g = 0.43, CI = [0.13, 0.90]).

### Other clinical, neuropsychological and sociodemographic data

We found that spatial suppression decreased with age in both groups (Figure 2B), a result reported for healthy participants in numerous studies (Betts et al. 2005; Betts, Sekuler, and Bennett 2009; Yazdani et al. 2015; Zhuang et al. 2017; Pitchaimuthu et al. 2017; Tadin et al. 2019; Deng et al. 2017; Karas and McKendrick 2012). The decrease was mostly explained by a decrease in motion sensitivity with age for small stimuli (Figure 2B).

We found that spatial suppression did not correlate with symptom severity assessed using BNSS (r = −0.14, p = 0.43), negative PANSS (r = 0.010, p = 0.96), positive PANSS (r = 0.21, p = 0.24), general PANSS (r = 0.19, p = 0.28) and total PANSS (r = 0.16, p = 0.38). Tadin and colleagues (Tadin et al. 2006) reported that spatial suppression did not correlate with symptom severity assessed by the Brief Psychiatric Rating Scale (BPRS) and the Scale for the Assessment of Positive Symptoms (SAPS), but correlated with the Scale for the Assessment of Negative Symptoms (SANS; r = −0.54, p = 0.03). Given the strong associations between SANS and negative PANSS (Rabany et al. 2011; van Erp et al. 2014; Norman et al. 1996; Kay, Opler, and Lindenmayer 1988; Peralta, Cuesta, and de Leon 1995), and SANS and BNSS (Kirkpatrick et al. 2011), it seems more likely that the apparent inconsistent findings are explained by a sampling error rather than by the use of different scales.

Next, we assessed the possible effect of medication on spatial suppression. We found a correlation of r(31) = 0.29 with p = 0.10 between spatial suppression and medication doses assessed using chlorpromazine equivalents. Tadin and colleagues (Tadin et al. 2006) found a correlation of r = −0.27 (personal communication by Duje Tadin). This evidence suggests that spatial suppression for motion is not mediated by medication. Furthermore, we found that spatial suppression did not correlate with tobacco consumption (number of cigarettes per month) in patients (r(31) = 0.047, p = 0.79) or in controls (r(29) = −0.14, p = 0.45).

Finally, we performed an exploratory data analysis to look at the relation between the perceptual variables and the overall functioning outcome using GAF. We found that in patients, but not in controls, spatial suppression decreased with GAF (Figure 2C). The interaction of GAF by group (F(1,60) = 3.88) had a p=0.053. The decrease was mostly explained by a strong decrease in motion sensitivity with GAF for small stimuli (Figure 2C). That is, the patients with worse functioning were the most sensitive to motion. A task for the future is to assess the robustness of these associations with GAF.

### Meta-analysis

To quantitatively assess the current evidence supporting a reduction of spatial suppression for motion in schizophrenia, we performed a meta-analysis that include the previous study (Tadin et al. 2006) and ours.The combined effect size was g = 0.47 (CI = [0.055, 0.88], p = 0.026, Figure 3A).

**Figure 3.**
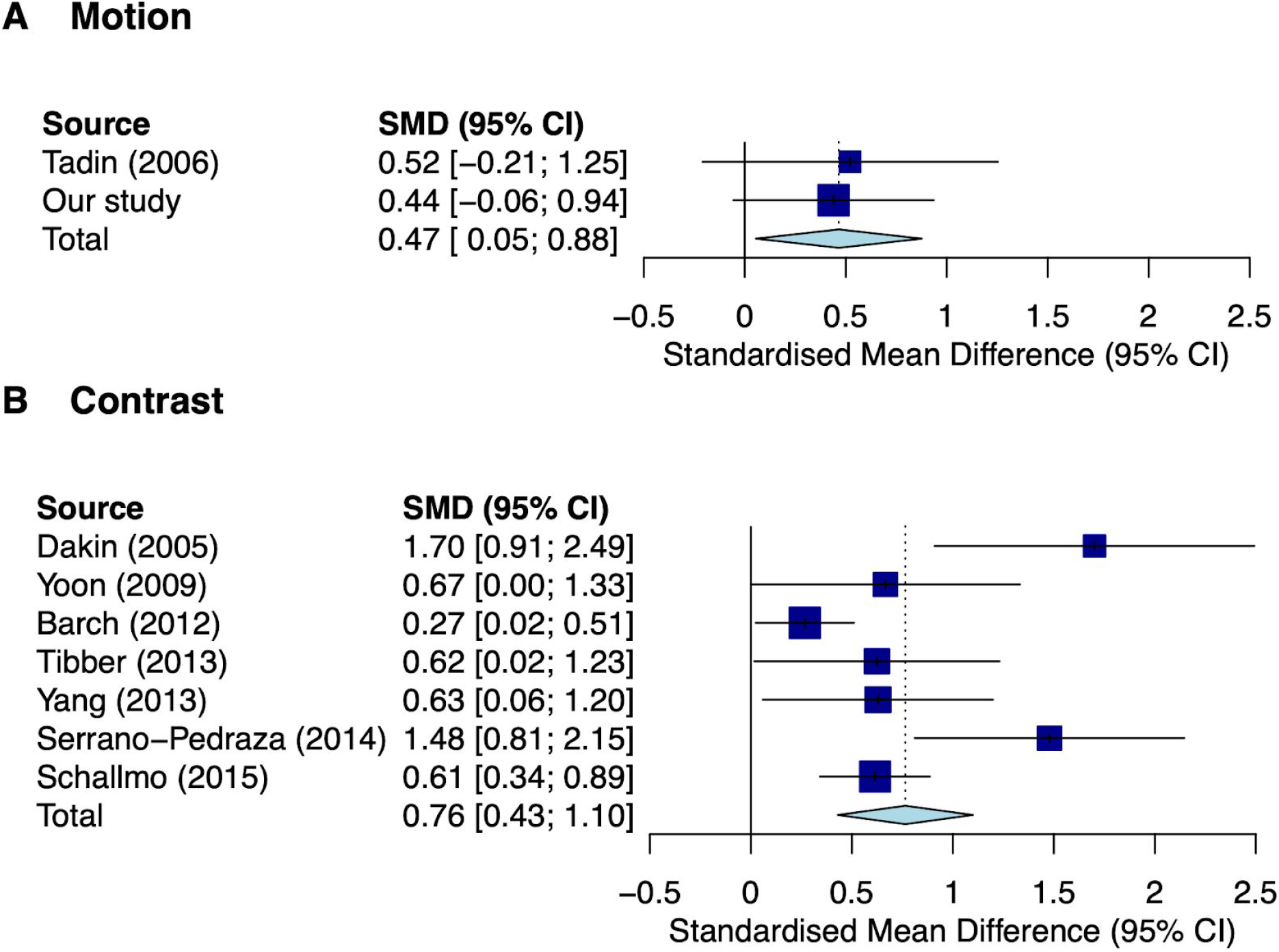
Random-effects model meta-analysis for the reduction of spatial suppression for motion (A) and contrast (B) in schizophrenia.

To compare the magnitude of the reduction of spatial suppression for motion in schizophrenia with that of contrast, we also performed a meta-analysis for contrast. We combined contrast suppression studies that measured appearance and performance (Kingdom and Prins 2016). For the appearance studies (Dakin, Carlin, and Hemsley 2005; Yang et al. 2013; Barch et al. 2012; Tibber et al. 2013; M-P Schallmo, Sponheim, and Olman 2015), the variable that we used to calculate the effect size was the reduction in perceived contrast—in the study of Schallmo and colleagues (M-P Schallmo, Sponheim, and Olman 2015) we used the condition in which the surround and the central stimuli had the same orientation as this is the condition that maximizes suppression (Solomon, Sperling, and Chubb 1993). For the performance studies (Serrano-Pedraza et al. 2014; Yoon et al. 2009), we used the suppression indexes reported in the studies—for both studies we used the condition in which the surround and the central stimuli had the same orientation (Yoon et al. 2009). For motion, the study of Tadin and our study measured performance and, as we described, the variable used was the suppression index.

The combined effect size for the reduction of contrast suppression in schizophrenia was g = 0.76 (CI = [0.43, 1.1], p = 7.5 x 10^-6^, Figure 3B). As there is evidence of publication bias (Egger’s test, intercept = 3.0, CI = [0.87, 5.2], p = 0.045), we recalculated the combined effect size using the method of Duval and Tweedie (2000). The trimmed and filled studies were Dakin (2005), Yoon (2009) and Serrano-Pedraza (2014). The recalculated effect size was g = 0.48 (CI = [0.13, 0.84], p = 0.0068), which suggests that the reduction of contrast suppression in schizophrenia cannot be just explained by publication bias. The effect size is, thus, similar for contrast and motion, although the evidence for contrast is larger because there are more studies having an effect in the same direction.

## DISCUSSION

We found evidence that patients with schizophrenia had a weaker spatial suppression than healthy controls—which is consistent with the study of Tadin and colleagues (Tadin et al. 2006)—and that the spatial suppression of patients, but not controls, increased with their IQ, which has not been previously described. Studies assessing the effect of IQ on spatial suppression in healthy participants had conflicting results, with two studies finding an increase in spatial suppression with IQ (Arranz-Paraíso and Serrano-Pedraza 2018; Melnick et al. 2013) and one showing no association (Troche et al. 2018). One could argue that because there is some overall evidence of an increase in spatial suppression with IQ, and patients with schizophrenia usually have lower IQ than healthy controls (Van Haren et al. 2019), a potential explanation for the weaker spatial suppression in patients with schizophrenia is their lower IQ. Our results are not consistent with this interpretation because in a subsample of 27 patients and 27 controls with matched IQ, the weaker spatial suppression in patients was maintained at the same level.

We found that spatial suppression decreased with age in healthy controls—an effect consistently found in previous studies (Betts et al. 2005; Betts, Sekuler, and Bennett 2009; Yazdani et al. 2015; Zhuang et al. 2017; Pitchaimuthu et al. 2017; Tadin et al. 2019; Deng et al. 2017; Karas and McKendrick 2012). Here, we found that this age-mediated perceptual alteration also occurs in patients with schizophrenia. In studies assessing visual cortex of monkeys, an age-related decrease in GABA-mediated inhibition was identified (Leventhal et al. 2003; Schmolesky et al. 2000), leading to the proposal that a reduction of GABAergic function plays a role in the decrease in spatial suppression with age (Betts et al. 2005). These findings, however, are different from those reported in humans. A recent study (Pitchaimuthu et al. 2017) using magnetic resonance spectroscopy (MRS) showed higher levels of GABA signal in the visual cortex of aged participants, in whom a decreased spatial suppression was confirmed. Interestingly, the same study found lower levels of glutamate in the same participants.

A decrease in GABA-mediated inhibition has also been proposed to explain the weaker spatial suppression for motion in schizophrenia (Tadin et al. 2006). However, pharmacological manipulations to increase the levels of GABA resulted in a decrease of spatial suppression in humans (Michael-Paul Schallmo et al. 2018) and did not affect the surround suppression of neurons in the monkey (Liu, Miller, and Pack 2018) linked to the perceptual spatial suppression (Liu, Haefner, and Pack 2016). Overall, current evidence suggests that a decline in GABAergic function cannot explain the perceptual differences in surround suppression between younger and aged participants, or between healthy controls and patients with schizophrenia.

Perceptual alterations in schizophrenia have also been associated with a glutamatergic hypofunction (Phillips and Silverstein 2013; Butler, Silverstein, and Dakin 2008; Yoon et al. 2013; Javitt and Sweet 2015), which is currently considered to play a central role in the pathogenesis of this disease (Uno and Coyle 2019; Moghaddam and Javitt 2012). Spatial suppression is functionally linked to gain control adjustments (Carandini and Heeger 2011), which are known to be mediated by glutamatergic neurotransmission (Daw, Stein, and Fox 1993; Butler, Silverstein, and Dakin 2008). For example, Kwon and colleagues (Kwon et al. 1992) showed that the response of neurons in the cat’s lateral geniculate nucleus to visual stimulation decreased by blocking NMDA glutamatergic receptors. Interestingly, they also showed that NMDAR blockers have a much larger effect on small stimuli than on large stimuli. This finding was observed using static stimulation, but is consistent with the weakened spatial suppression that we found in patients with schizophrenia, which was mostly driven by their poor motion sensitivity for small stimuli.

Apart form the decrease in spatial suppression, we found that patients also performed the motion discrimination task less accurately than healthy controls. Tadin and colleagues (Tadin et al. 2006) found evidence, albeit weaker, in the same direction. This deficit adds to other deficits in motion perception tasks found in patients with schizophrenia such as discriminating the motion direction of signals distributed across space (Y. Chen 2011; Carter et al. 2017) or the difference in speed of two stimuli (Yue Chen et al. 1999). In contrast with surround suppression, changes in accuracy can be more easily interpreted as being caused by a generalized behavioural deficit associated with the condition, such as a lack of attention or motivation. To assess the generalized deficit, in this study we included trials in which the stimuli should be easy to discriminate (Dakin, Carlin, and Hemsley 2005; Yoon et al. 2013; Barch et al. 2012). Consistent with patients losing attention or motivation more frequently than controls, we found that patients were less accurate in easy trials—had more lapses. This is a known deficit for other behavioural tasks not assessing perceptual sensitivity (Cornblatt and Keilp 1994). Only lapses, however, could not explain the worse performance of patients in the task. Our measure of sensitivity, which is not contaminated by lapses (Prins 2013), indicates that patients had a genuine worse motion sensitivity than controls. This finding might be also consistent with a glutamatergic hypofunction origin of schizophrenia as a recent MRS study showed that healthy participants with decreased glutamate levels in a visual motion area performed worse in a motion discrimination task similar to the one used in this study (Michael-Paul Schallmo et al. 2019).

At present, it is unclear whether the reduced sensitivity and spatial suppression that we found is associated with processes occurring proximally to the etiology of psychosis or rather associated to the chronicity of the illness. This ambiguity could be resolved by measuring motion sensitivity and spatial suppression in patients with a first episode of psychosis.

Surround suppression is a fundamental sensory process with well-studied physiological mechanisms and functions (Carandini and Heeger 2011; Butler, Silverstein, and Dakin 2008), which has been reported to be compromised in schizophrenia. For the perception of contrast, numerous studies have found weakened surround suppression (Barch et al. 2012; Yang et al. 2013; Tibber et al. 2013; Serrano-Pedraza et al. 2014; Yoon et al. 2009; M-P Schallmo, Sponheim, and Olman 2015; Dakin, Carlin, and Hemsley 2005)) with a modest effect size (0.48 after correcting for publication bias) according to our meta-analysis. For the perception of motion, only our study and the study of Tadin and colleagues (Tadin et al. 2006) have been reported. Both studies (our study controlling for IQ) suggest a weakened surround suppression in schizophrenia with a combined effect size (0.47) similar to the effect size for contrast. We think that further studies are needed to establish how robust the effect is. Identifying robust perceptual alterations in schizophrenia is important as they might point towards neural dysfunctions such as imbalances of certain neurotransmitters (Butler, Silverstein, and Dakin 2008; Javitt and Sweet 2015; Yoon et al. 2013; Phillips and Silverstein 2013) and they can provide much needed objective measures of disease severity and treatment outcome.

## ACKNOWLEDGMENTS

This work was supported by the Fundación Alicia Koplowitz, Fundació CELLEX, La Caixa Foundation Health Research, CIBERSAM, CIBERER (Refs: 15/00010), the Generalitat de Catalunya (PERIS SLT002/16/00338, SLT006/17/00362, 2014SGR1265, 2017SGR1355, SLT006/17/00345), the Spanish Ministry of Science, Innovation and Universities and European Regional Development Fund (Refs: BFU2015-65315-R, RTI2018-094190-B-I00, PI08/0208, PI11/00325, PI14/00612), Instituto Carlos III/FEDER (Refs: FIS 17/00234, PIE 16/00014) and by the CERCA Programme/Generalitat de Catalunya. Part of this work was developed at the building Centro Esther Koplowitz, Barcelona. We thank Deanna Barch, Steve Dakin, Ariel Rokem, Michael-Paul Schallmo, Duje Tadin and Ignacio Serrano-Pedraza for sharing data or statistics from their studies and Heike Stein for comments on the manuscript.

## Notes

### Competing Interest Statement

The authors have declared no competing interest.

